# When is warmer better? Disentangling within- and between-generation effects of thermal history on survival

**DOI:** 10.1101/2023.02.13.528408

**Authors:** Adriana P. Rebolledo, Carla M. Sgrò, Keyne Monro

## Abstract

1. Understanding the fitness consequences of thermal history is necessary to predict organismal responses to global warming. This is especially challenging for ectotherms with complex life cycles, since distinct life stages can differ in thermal sensitivity, acclimate to different thermal environments, and accrue responses to acclimation within and between generations.
2. Although acclimation is widely hypothesized to benefit organisms by helping them (or their offspring) to compensate for negative impacts of environmental change, equivocal support for this hypothesis highlights the need to assess alternatives. However, assessments that do so in ways that explicitly dissect responses across life stages and generations remain limited.
3. We assess alternative hypotheses for acclimation responses (none, beneficial, colder-is-better, and warmer-is-better) within and between generations of an externally-fertilizing marine tubeworm whose vulnerability to warming rests on survival at early planktonic stages (gametes, embryos, and larvae). We start by acclimating parents, gametes, and embryos to ambient and projected warmer temperatures (17 °C and 22 °C) factorially by life stage. We then rear individuals with differing acclimation histories to the end of larval development at test temperatures from 10 °C to 28 °C (upper and lower survival limits) to estimate thermal survival curves for development, and compare curves among acclimation histories.
4. We show that survival curves are most responsive to parental acclimation followed by acclimation at embryogenesis, but are buffered against acclimation at fertilization. Moreover, curves respond independently to acclimation within and between generations, and respond largely as predicted by the warmer-is-better hypothesis, despite the semblance of beneficial acclimation after successive acclimations to warmer temperature.
5. Our study demonstrates the varied nature of thermal acclimation, and the importance of considering how acclimation responses aggregate across complex life cycles when predicting vulnerability to warming.

## Introduction

Rising temperatures from global warming are affecting biological function from molecules to ecosystems, with key implications for ectotherms that dominate the globe and use environmental temperature to thermoregulate to varying degrees (Deutsch *et al.* 2008; Seebacher, White & Franklin 2015; Clarke 2017). Population persistence rests on the capacity to maintain Darwinian fitness (survival and reproduction) in the face of warming, spurring efforts to identify thermally-sensitive life stages acting as lifecycle bottlenecks (Dahlke *et al.* 2020; Sunday 2020) and the extent to which thermal history — the temperatures experienced by prior generations or life stages — buffers them against warming or worsens its impacts (Kellermann, van Heerwaarden & Sgrò 2017; Donelson *et al.* 2018). Recent work points to greater thermal sensitivity at early life stages relative to juveniles and adults (Pandori & Sorte 2019; Dahlke *et al.* 2020; Kingsolver & Buckley 2020; Taylor *et al.* 2021), yet the role of thermal history remains unclear. Despite mounting evidence of its downstream impacts on thermal sensitivity within and between generations (e.g., Huey *et al.* 1995; Zamudio, Huey & Crill 1995; Crill, Huey & Gilchrist 1996; Steigenga & Fischer 2007; Le Roy & Seebacher 2018; Cavieres *et al.* 2019; Diaz *et al.* 2021), there is a need to explicitly dissect those effects across the complex lifecycles of ectotherms, especially at early life stages that may define vulnerability to warming.

It has long been proposed that prior acclimation to thermal stress may enhance the fitness of organisms who re-encounter that stress later in life, giving them an adaptive advantage over non-acclimated organisms (Levins 1969; Hochachka & Somero 1984; Hoffmann & Parsons 1991). This so-called beneficial acclimation hypothesis (Leroi, Bennett & Lenski 1994) has been framed more broadly in the context of adaptive phenotypic plasticity to other environmental stressors (Via *et al.* 1995; Ghalambor *et al.* 2007). It has also been extended to parental acclimation, positing that parental exposure to environmental stress (thermal or otherwise) primes offspring to tolerate that stress better than offspring from non-stressed parents (Agrawal, Laforsch & Tollrian 1999; Galloway & Etterson 2007; Donelson *et al.* 2012). Despite support from individual case studies, however, multiple surveys of the empirical literature now challenge the generality of the beneficial acclimation hypothesis in its various forms. Regardless of whether acclimation responses are surveyed within generations (Woods & Harrison 2002; Angilletta 2009; Sgrò, Terblanche & Hoffmann 2016) or across them (Uller, Nakagawa & English 2013; Sgrò, Terblanche & Hoffmann 2016; Donelson *et al.* 2018; Sánchez□Tójar *et al.* 2020), acclimation seems to be detrimental just as often as it is beneficial, especially for fitness-related traits tested after acclimation to stressful conditions (Sánchez□Tójar et al. 2020). Responses to acclimation may therefore be more complex and varied than predicted by the beneficial acclimation hypothesis, highlighting the need to jointly consider alternative hypotheses that make different predictions.

An expanded perspective is especially important when considering the adaptive value of thermal acclimation. Given the nonlinear, unimodal relationship between temperature and fitness, thermal performance curves (Figure 1) can respond to acclimation in ways that affect whether and where they cross, thereby changing the rank order of fitness, in the range of test temperatures (Huey and Berrigan 1996). Without an idea of curve shapes, and where fitness is measured in relation to features such as limits and optima, responses to acclimation may be misconstrued (Angilletta 2009; Schulte, Healy & Fangue 2011). Accordingly, Huey and Berrigan (1996) and Huey *et al.* (1999) derived a set of competing hypotheses of acclimation that predict different responses of performance curves. If acclimation is beneficial, compensating for a stressfully warmer temperature that reduces fitness otherwise, then the thermal optimum should increase at no cost to maximal fitness (Figure 1A). Conversely, if warm acclimation is detrimental, then non-stressed ‘colder’ organisms should do better than ‘warmer’ stressed ones (Figure 1B). If thermodynamic effects (higher reaction kinetics at warmer temperatures) scale up to fitness, however, then warmer is better (Huey & Kingsolver 1989; Angilletta, Huey & Frazier 2010) — in other words, warm-acclimated organisms should have higher thermal optima, and do better at those optima, than non-acclimated organisms with lower optima (Figure 1C), at least until upper thermal limits are met. Last, no acclimation effect (Figure 1D) might reflect past selection for developmental mechanisms that buffer fitness against perturbation (Huey *et al.* 1999). Acclimation may of course have more nuanced effects, and other hypotheses relating to curve position and breadth have also been proposed (e.g., Izem & Kingsolver 2005). Nevertheless, those above offer a key framework for characterizing acclimation that has been applied to within-generation responses (e.g., Treasure & Chown 2019), but rarely to responses passed from one generation to the next.

**Figure 1A-D.**
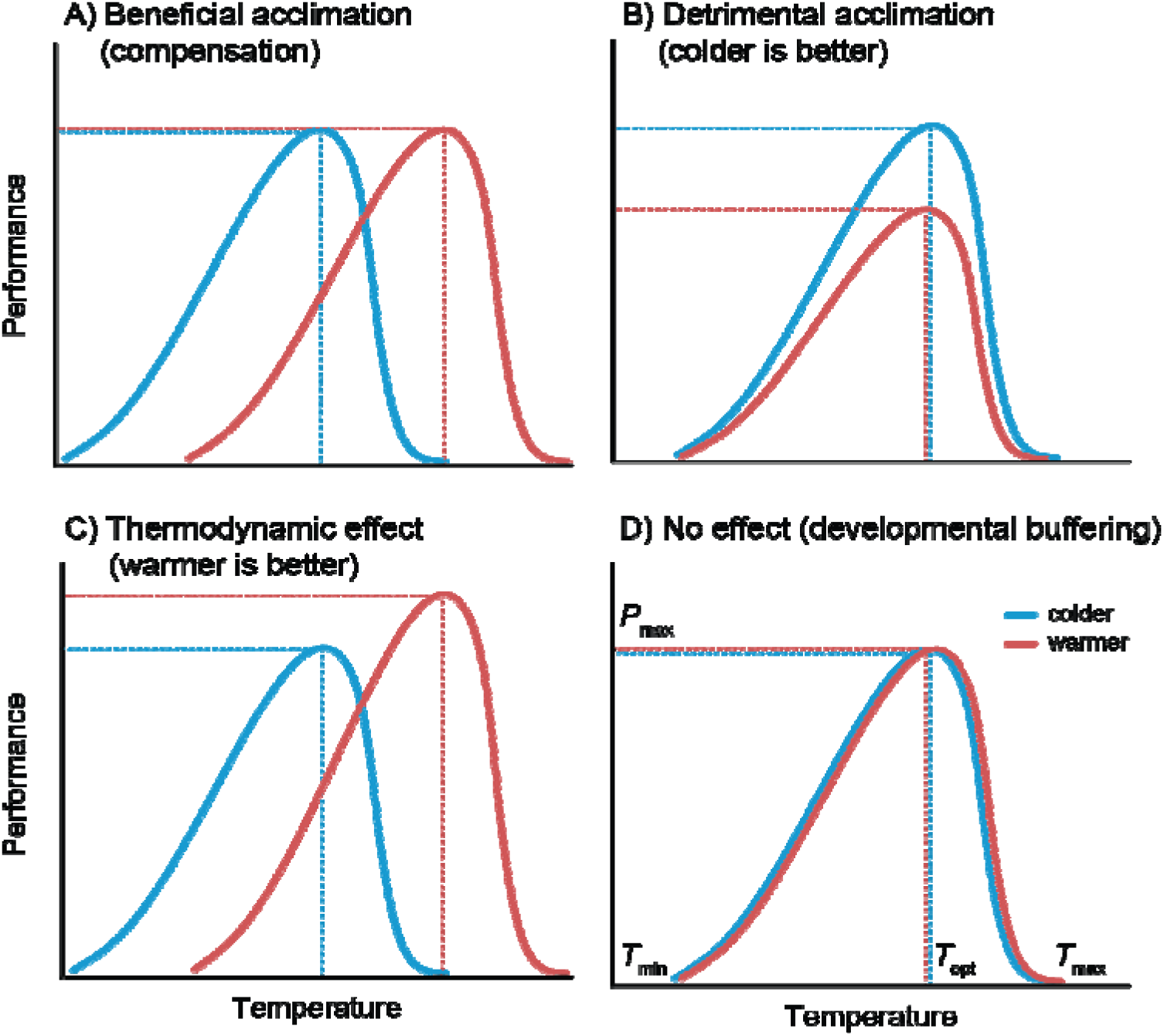
Alternative hypotheses of acclimation and their predictions for thermal performance (adapted from Huey *et al.* 1999). Performance is a unimodal function of temperature, rising from zero at its lower thermal limit (*T*_min_) to a maximum (*P*_max_) at its thermal optimum (*T*_opt_), then returning to zero at its upper thermal limit (*T*_max_). A) If beneficial acclimation compensates for a stressfully warmer temperature, then the thermal optimum should increase at no cost to maximal performance. B) If warm acclimation is detrimental, then non-stressed ‘colder’ organisms should do better overall than ‘warmer’ stressed ones. C) If warmer is better due to thermodynamic effects, then warm-acclimated organisms should have a higher thermal optimum, and do better at that optimum, compared to non-acclimated organisms with lower optima. D) No effect of acclimation may support developmental mechanisms that buffer performance against perturbation.

Uncertainty surrounds the relative strengths of responses to acclimation within *versus* between generations, and their aggregation across complex lifecycles (Sgrò, Terblanche & Hoffmann 2016; Williams *et al.* 2016; Donelson *et al.* 2018). Ideally, acclimation should let organisms shift their thermal optima to match each successive temperature they face, although thermodynamic effects can favor optima above ambient temperatures (Asbury & Angilletta 2010). However, theory emphasizes the role of thermal variability (how well temperature cues at one life stage predict selection at another) in moderating such responses (Angilletta 2009; Leimar & McNamara 2015), and predictions rest on whether performance aggregates over time multiplicatively through effects on survival, or additively through effects on reproduction (Buckley & Kingsolver 2021). For example, increases in curve breadth may be favored over shifts in optima by variability within generations if performance is multiplicative, or by variability among generations if performance is additive (Lynch & Gabriel 1987; Gabriel & Lynch 1992; Gilchrist 1995). Limited empirical support for such predictions nevertheless suggests that unpredictable cues also lead to other responses such as bet-hedging (Buckley & Kingsolver 2021; Seebacher & Little 2021), or that benefits of acclimation are countered by trade-offs — for example, if responses that initially buffer against thermal stress prove detrimental later in life (Loeschcke & Hoffmann 2002; Angilletta 2009). Ultimately, studies are needed that not only test alternative hypothesis of acclimation, but also disentangle responses within and across generations. To date, such studies are rare and largely focus on insects (Huey *et al.* 1995; Crill, Huey & Gilchrist 1996; Steigenga & Fischer 2007; but see Kielland, Bech & Einum 2017, or Le Roy & Seebacher 2018).

Here, we compare thermal survival curves after a factorial manipulation of thermal history to assess alternative hypotheses of acclimation, and disentangle responses within and between generations, in the externally-fertilizing tubeworm, *Galeolaria caespitosa.* Like most aquatic ectotherms, its vulnerability to warming rests on survival at early planktonic stages (gametes, embryos, and larvae) dispersed passively by currents (Walsh *et al.* 2019; Byrne *et al.* 2020; Dahlke *et al.* 2020), potentially decoupling the temperatures experienced in early life from parental temperatures (Lotterhos, Albecker & Trussell 2021). Parents can still modify offspring responses to temperature by loading protective proteins into gametes, or passing on physiological damage (Hamdoun & Epel 2007; Guillaume, Monro & Marshall 2016; Chirgwin *et al.* 2018), and temperatures at fertilization and embryogenesis can further influence later responses to temperature (Chirgwin, Connallon & Monro 2021; Rebolledo, Sgrò & Monro 2021). This biology therefore presents rare scope to dissect cumulative responses to acclimation across key stages of the life cycle that govern adult abundances and dynamics. Using a split-cohort design to standardize genetic backgrounds across stages, we start by acclimating parents, gametes, and embryos to ambient and projected warmer temperatures (17 °C and 22 °C) factorially by life stage. We then rear individuals with differing acclimation histories to the end of larval development at test temperatures from 10 °C to 28 °C (upper and lower survival limits) to estimate thermal survival curves for development and assess how they respond to acclimation within and between generations.

## Materials and methods

### Study species and sampling

*Galeolaria caespitosa* (henceforth *Galeolaria*) is a calcareous tubeworm native to rocky shores of southeastern Australia, where it acts as an ecosystem engineer by forming dense colonies of tubes used as habitat and refuge by associated communities (Wright & Gribben 2017). Sessile adults breed year-round by releasing gametes into the sea for external fertilization (Chirgwin, Marshall & Monro 2020). Embryos develop into functionally-independent larvae ~24 hours later, then larvae develop in the plankton for another ~2-3 weeks until rapid changes in size, morphology, and behavior signal the end of planktonic life (readiness to settle and recruit into sessile populations; Marsden & Anderson 1981). These early stages are thermal bottlenecks in the life cycles of external fertilizers (Byrne 2011; Walsh *et al.* 2019; Dahlke *et al.* 2020), and effects of thermal history can persist from one stage to the next (Chirgwin, Connallon & Monro 2021; Rebolledo, Sgrò & Monro 2021), but the aggregate effects on early survival are unknown.

We sampled cohorts of parents between June and December 2020 from a natural population at Brighton, Port Phillip Bay, Victoria, where water temperatures range from 9–24 °C throughout the year (Chirgwin *et al.* 2018). The region is a marine hotspot that is warming much faster than the global average rate (Hobday & Pecl 2014), is projected to warm ~2–4 °C by the century’s end (RCP8.5 relative to 1980-1999; Lough, Sen Gupta & Hobday 2012), and is prone to heatwaves in excess of average warming (Oliver *et al.* 2017). Each cohort of parents was transferred in seawater to Monash University, held for several hours at the natural temperature to minimize transfer stress, then divided among replicate tanks of fresh aerated seawater and gradually adjusted to the designated parental temperature (see below).

### Experimental overview

We factorially manipulated parental temperature (17 °C *versus* 22 °C), temperature at fertilization (17 °C *versus* 22 °C), and temperature at embryogenesis (17 °C *versus* 22 °C), then estimated thermal survival curves for larval development (see Figure 2). Prior acclimation temperatures are the mean annual and summer temperatures in nature, with the latter projected to occur more frequently in coming years (Lough, Sen Gupta & Hobday 2012; Chirgwin *et al.* 2017), and include the thermal optima for fertilization and embryogenesis (Rebolledo, Sgrò & Monro 2020). Thermal survival curves were based on eight test temperatures that span the lower and upper survival limits (10–28°C) for larval development (Rebolledo, Sgrò & Monro 2020).

**Figure 2.**
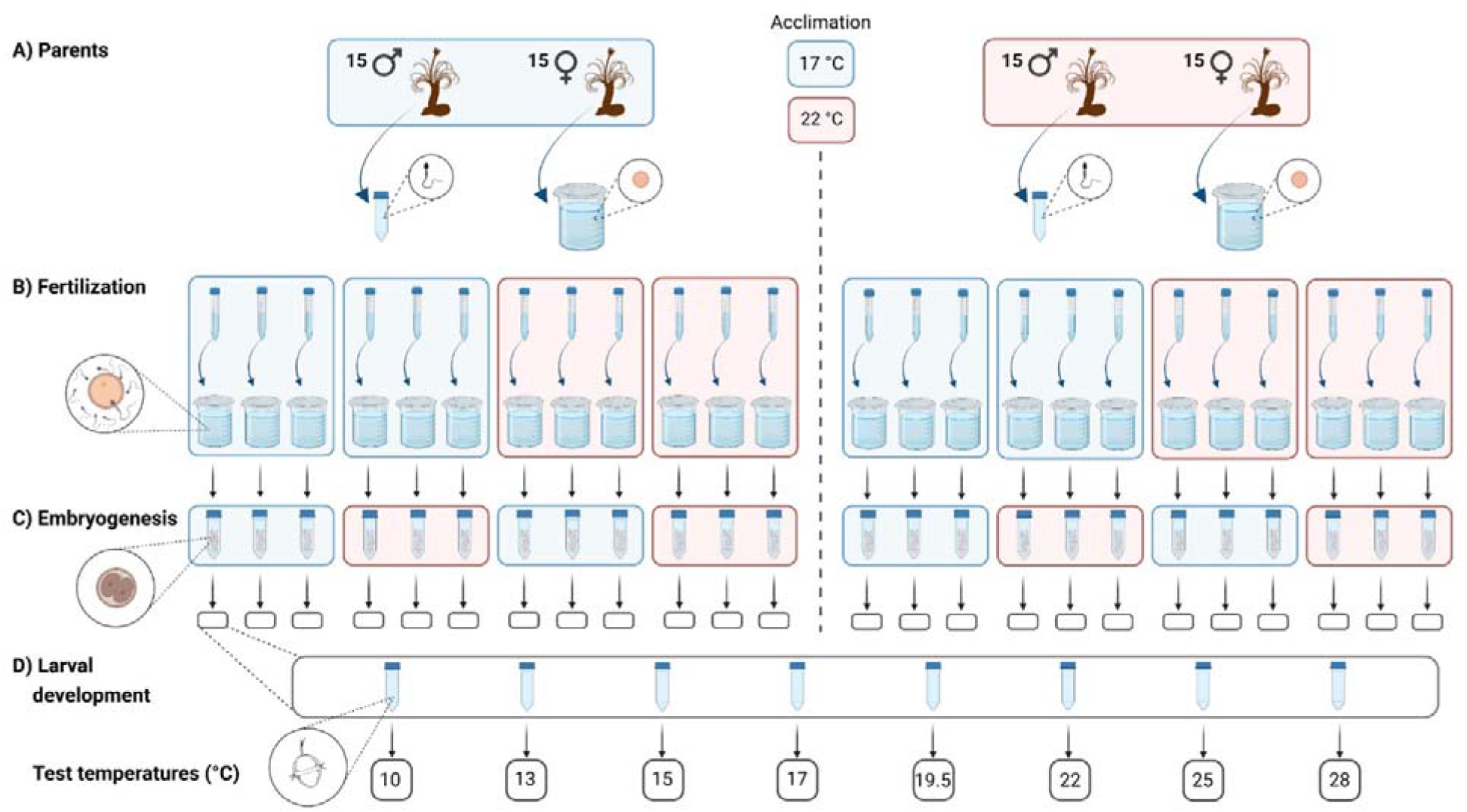
Factorial manipulation of thermal history. A) Each cohort of parents was acclimated at either 17°C or 22°C for one month, then gametes were collected from fifteen males and fifteen females per temperature and pooled by sex. B) Gametes from each parental temperature were used to initiate replicate fertilizations at 17 and 22 °C, so that fertilization and parental temperatures were crossed factorially and had six replicate vials per factorial combination. After 30 minutes of gamete contact, vial contents were rinsed to remove sperm and re-suspended in fresh seawater at the same temperature until zygotes divided into two cells (1–2 hours later). C) Samples of the resulting embryos per vial were transferred to new vials at 17°C and 22 °C, so that temperatures at this stage and prior stages were crossed factorially and had three replicate vials per factorial combination, until completing embryogenesis ~24 hours later. D) Thirty resulting larvae per vial were randomly allocated to each of eight test temperatures (10 °C to 28 °C), so that temperatures at this stage and prior stages were again crossed factorially and had three replicate vials per factorial combination. Larvae were fed *ad libitum* and monitored throughout development (up to three weeks depending on temperature) until they had either died or successfully completed development. Figure created with Biorender.

Except for parental environment (see below), thermal environments were manipulated, and survival assayed, in replicate vials of filtered, pasteurized seawater (loosely capped for oxygen flow) suspended upright in water baths held at designated temperatures (± 0.1 C) by controlled immersion heaters (Grant Optima TX150). Vials in survival assays each contained thirty individuals monitored throughout larval development. For each cohort of parents, replicates were generated in an incomplete block design with larval temperatures assigned haphazardly to blocks. Each block comprised three replicates per factorial combination of acclimation temperatures, assayed at five to eight larval temperatures (it was not logistically feasible to assay all eight larval temperatures per block). Hence, all replicates per block were derived from subsets of material from the same parents, assayed for survival concurrently under identical conditions aside from the factorial manipulation of temperature. This design was then replicated for each of four cohorts of parents. Ultimately, there were 574 vials (2 parental temperatures x 2 fertilization temperatures x 2 temperatures at embryogenesis x ~6 larval temperatures on average x 4 blocks x 3 replicates per block, less 2 replicates lost to contamination), with survival scored for ~17000 individuals grouped by vial (Figure 2).

### Manipulation of parental temperature

Since gametogenesis is continuous and gametes can ripen in under two weeks, each cohort of parents was acclimated for one month at either 17°C or 22°C prior to the steps below. Cohorts were acclimated in replicate tanks per temperature, fed a mix of live microalgae *ad libitum* every second day, and had seawater replaced weekly.

### Gamete collection and manipulation of temperature at fertilization

For each cohort of parents, gametes were collected from fifteen males and fifteen females per parental temperature (to minimize the effects of genetic incompatibilities; Marshall & Evans 2005; Chirgwin *et al.* 2017), with sexes chosen from different tanks. Each adult was extracted from its tube and placed in a dish with ~1 mL of fresh filtered seawater at 17 °C to spawn. Gametes were immediately collected, checked for quality based on appearance of eggs and motility of sperm, then pooled by sex and used within the hour before viability declines (Rebolledo, Sgrò & Monro 2020). Pooled eggs were diluted to ~250 cells mL^-1^ before use. Pooled sperm were kept concentrated at ~10^7^ cells mL^-1^ to minimize activity-dependent aging before use (Kupriyanova 2006; Chirgwin, Marshall & Monro 2020).

Gametes from each parental temperature were used to initiate replicate fertilizations at 17 and 22 °C, so that fertilization and parental temperatures were crossed factorially and had six replicate vials per factorial combination (Figure 2). To initiate each fertilization, 4.5 mL of pooled eggs and 0.5 mL of pooled sperm were separately adjusted to the required temperature over 30 min, then combined at that temperature. After 30 min of gamete contact (which maximizes fertilization success across the temperatures here; Rebolledo, Sgrò & Monro 2020), the contents of each vial were rinsed through 0.25μm mesh with seawater to remove sperm, then re-suspended in 45 mL of fresh seawater until zygotes divided into two cells (the earliest that they could be distinguished from unfertilized eggs under a stereomicroscope) ~1–2 hours later depending on temperature.

### Manipulation of temperature at embryogenesis

Samples of the resulting embryos per vial were transferred to new vials at 17°C and 22 °C, so that temperatures at this stage and prior stages were crossed factorially and had three replicate vials per factorial combination (Figure 2). All embryos were at a similar point in development (two cells) when exposed to acclimation temperatures. Embryos were maintained in 45 mL of seawater, sufficient to avoid oxygen-limitation (Chirgwin *et al.* 2018) until they completed development into actively swimming, feeding larvae ~24 hours later.

### Assaying survival of larval development

Thirty resulting larvae per vial were randomly allocated to each of 3-4 replicate vials per test temperature (10, 13, 15, 17, 19.5, 22, 25, or 28 °C) so that temperatures at this stage and prior stages were again crossed factorially (Figure 2). Larvae were maintained in 10 mL of seawater, sufficient to avoid oxygen-limitation (Chirgwin *et al.* 2018), and fed a mix of non-live microalgae *ad libitum* (~1 x 10^4^ cells mL^-1^ every second day, with seawater partially replaced at this point). We used non-live microalgae to avoid potential confounding of food availability if live microalgae differed in growth rate at different temperatures. After the first week, all but one of the vials per test temperature were sampled daily to check developmental progression (which is incomplete before this time; Rebolledo, Sgrò & Monro 2020), leaving the remaining vial undisturbed. Sampling continued for up to three weeks depending on temperature, ending when all larvae in the sampled vials had either died or successfully completed development. At this point, survival was assayed in the remaining undisturbed vial (vials used to check developmental progression did not contribute data to the final analysis).

### Modelling thermal survival curves

We fitted thermal survival curves to binary data (scores of 1 if individuals survived development and 0 otherwise) using a binomial mixed-effects regression model fitted with Laplace approximation in the *lme4* package (version 1.1-26; Bates *et al.* 2015) for *R* 4.0.5 (R Core Team, 2021). This model gives the same results as one fitted to the proportions of survivors per vial (i.e., 574 rows of data, each representing one vial), but was preferred here because it performed better in checks of model assumptions. Based on shapes of unconstrained smoothers fitted to data, survival curves were modelled as cubic functions of test temperature using orthogonal polynomials. Acclimation temperatures and all possible interactions with test temperature were initially included as fixed effects, before three- and four-way interactions were excluded to avoid overfitting (doing so did not reduce model fit: joint *χ^2^* test = 2.06, d.f. = 13, *P* = 0.99). Block was also included as a fixed effect, as was vial as a random effect. Checks of model assumptions using the *DHARMa* package (version 0.4.1; Hartig 2021) showed no violations. Fixed effects were tested using Wald *X*^2^ tests (Bolker *et al.* 2009) in the *car* package (version 3.0-10; Fox & Weisberg 2019). Pairwise contrasts were done for significant effects using *z* tests of log odds ratios (Sidak-adjusted as necessary) in the *emmeans* package (version 1.6.0; Lenth *et al.* 2021).

### Estimates and confidence intervals of curve descriptors

For acclimation temperatures with significant effects on survival curves, we extracted curve descriptors from the fitted model. Thermal optimum (*T*_opt_) was calculated as the temperature of maximal survival (*P*_max_), and thermal limits (*T*_min_ and *T*_max_) were calculated as the lower and upper temperatures at which survival was 5% of its maximum. We took this approach because binary data may approach 0% via an asymptote, thereby limiting the biological meaning of *T*_min_ and *T*_max_ at complete mortality (Kellermann *et al.* 2019), and results were qualitatively unchanged when they were calculated at complete mortality. We also explored thermal breadth (calculated as the temperature range at which survival was at least 50% of its maximum; Sinclair *et al.* 2016) and thermal tolerance (calculated as *T*_max_-*T*_min_), but such descriptors did not add to the conclusions drawn from other results so are not presented here.

To compare curve descriptors between acclimation temperatures, and thereby evaluate hypotheses in Figure 1, we used parametric bootstrapping (implemented in the *boot* package version v1.3-27; Canty & Ripley 2021) to estimate the mean and 95% confidence interval of each descriptor based on 1000 bootstrap replicates of the fitted model. Descriptors were considered to differ significantly between temperatures if their 95% confidence intervals did not overlap. Because this may be a conservative indicator of significance, we also calculated means and 95% confidence intervals for contrasts between temperatures (in this case, descriptors are significantly different if the confidence interval for their contrast excludes 0). Both indicators of significance gave similar results, so we rely on the former here.

## Results

### Modelling thermal survival curves

Parental temperature and temperature at embryogenesis interactively affected mean survival regardless of test temperature (Table 1 and Figure 3), but independently affected thermal survival curves — that is, mean survival at different test temperatures (Table 1 and Figure 4).

**Table 1.** Effects of parental temperature, temperature at fertilization, and temperature at embryogenesis on the probability of surviving larval development at different test temperatures. Survival was modeled as a cubic function of test temperature in a binomial mixed-effects regression. Higher-order interactions were non-significant (joint χ test = 2.06, d.f. = 13, *P* = 0.99) and were removed from the model to avoid overfitting. *P*-values in bold are significant at *a* = 0.05.

| Fixed effects | $\chi^2$ | d.f. | $P$ |
| --- | --- | --- | --- |
| Parental temperature | 7.36 | 1 | **<0.01** |
| Temperature at fertilization | 0.30 | 1 | 0.58 |
| Temperature at embryogenesis | 0.00 | 1 | 0.96 |
| Test temperature for larvae (linear, quadratic, and cubic trends) | 4970.21 | 3 | **<0.001** |
| Parental temperature x temperature at fertilization | 0.14 | 1 | 0.70 |
| Parental temperature x temperature at embryogenesis | 3.89 | 1 | **0.05** |
| Temperature at fertilization x temperature at embryogenesis | 0.01 | 1 | 0.94 |
| Parental temperature x test temperature | 120.20 | 3 | **<0.001** |
| Temperature at fertilization x test temperature for larvae | 1.97 | 3 | 0.58 |
| Temperature at embryogenesis x test temperature for larvae | 31.25 | 3 | **<0.001** |
| Block | 35.50 | 3 | **<0.001** |

**Figure 3.**
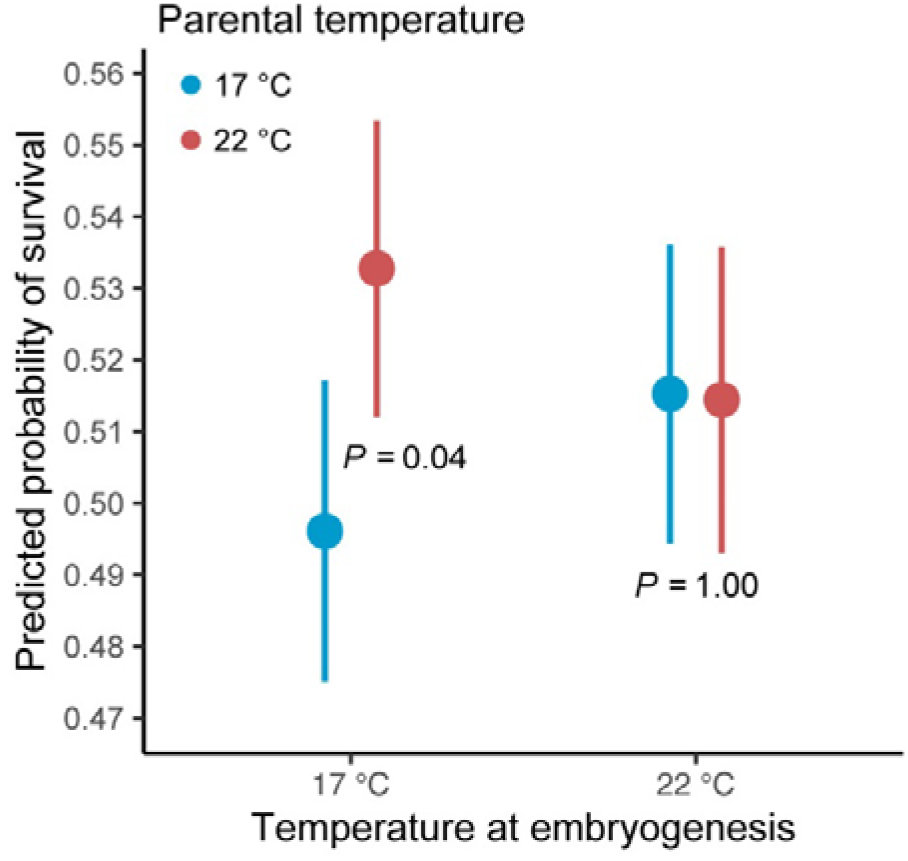
Interactive effects of parental temperature and temperature at embryogenesis (17 °C in blue *versus* 22 °C in red) on the predicted probability of surviving larval development, regardless of test temperature. Solid points are means (± 95% confidence intervals). Parental acclimation at 22 °C improved survival when followed by embryogenesis at 17 °C but not when followed by embryogenesis at 22 °C (see test statistics in text). Survival was unaffected by temperature at fertilization.

**Figure 4.**
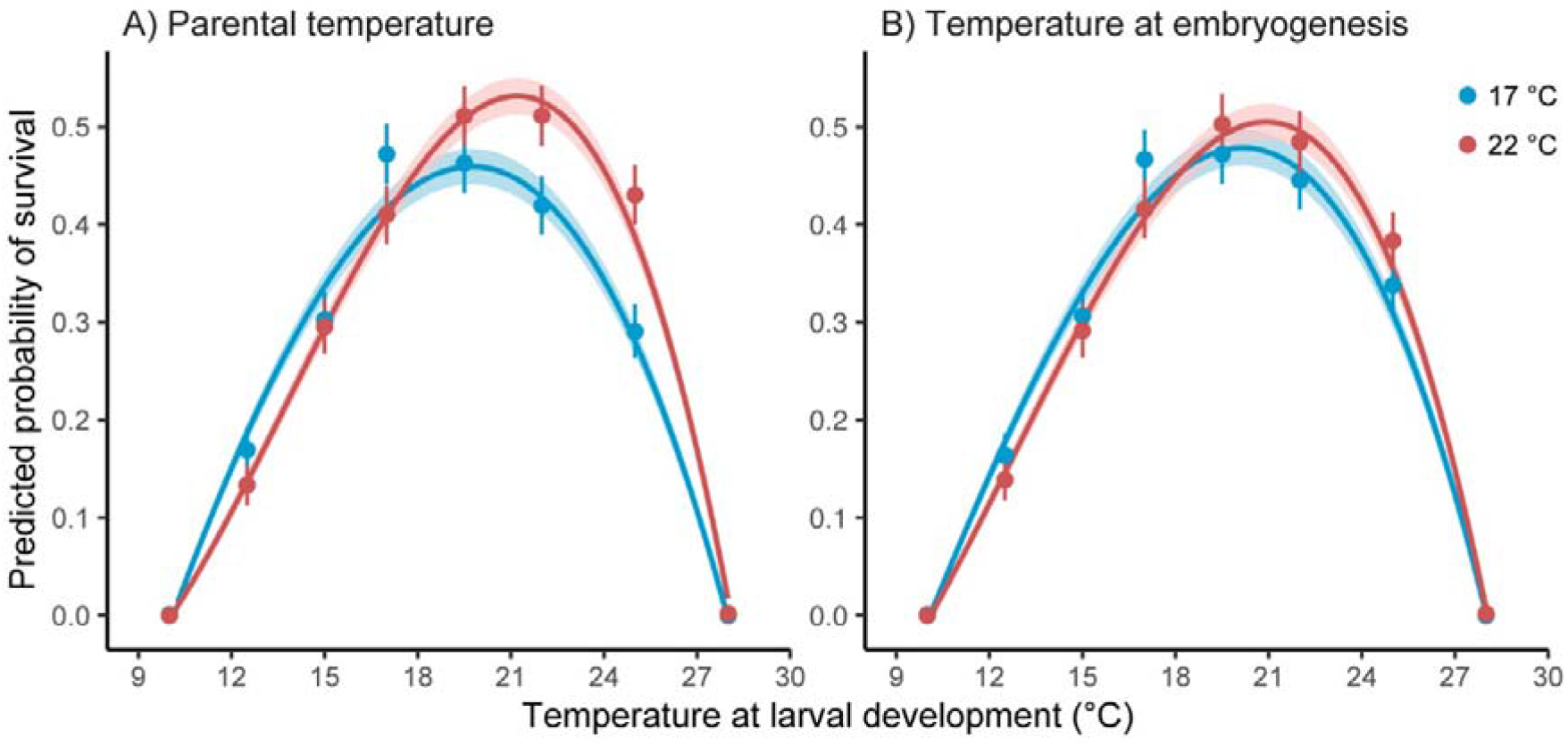
Independent effects of A) parental temperature and B) temperature at embryogenesis (17 °C in blue *versus* 22 °C in red) on the predicted probability of surviving larval development at different test temperatures. Survival curves for parental temperatures are averaged across temperatures at embryogenesis, and *vice versa.* Shaded areas are 95% confidence intervals of curve predictions. Points are observed proportions (± 95% confidence intervals) of survival at each test temperature. Survival was unaffected by temperature at fertilization.

The interactive effect of acclimation temperatures on survival was detected because parental acclimation at 22 °C increased the probability of survival when embryogenesis occurred at 17 °C (*z* = |2.54|, *P* = 0.04), but not when it occurred at 22 °C (*z* = |0.06|, *P* = 1.00) (Figure 3). The independent effects of acclimation temperatures on thermal survival curves were detected because linear trends (average slopes of curves in Figure 4) were consistently positive (Figure 5A), but significantly more so after parental acclimation at 22°C (*z* = |9.53|, *P* < 0.01), and after embryogenesis at 22°C (*z* = |5.01|, *P* < 0.01). Likewise, cubic trends (initial slopes of curves in Figure 4) were consistently negative (Figure 5C), but significantly more so after parental acclimation at 22°C (*z* = |6.16|, *P* < 0.01; Figure 5C), and after embryogenesis at 22°C (*z* = |3.31|, *P* < 0.01). Both linear and cubic trends were more sensitive to parental temperature than to temperature at embryogenesis. However, quadratic trends (breadths of curves in Figure 4) were unresponsive to either acclimation temperature (Figure 5B).

**Figure 5.**
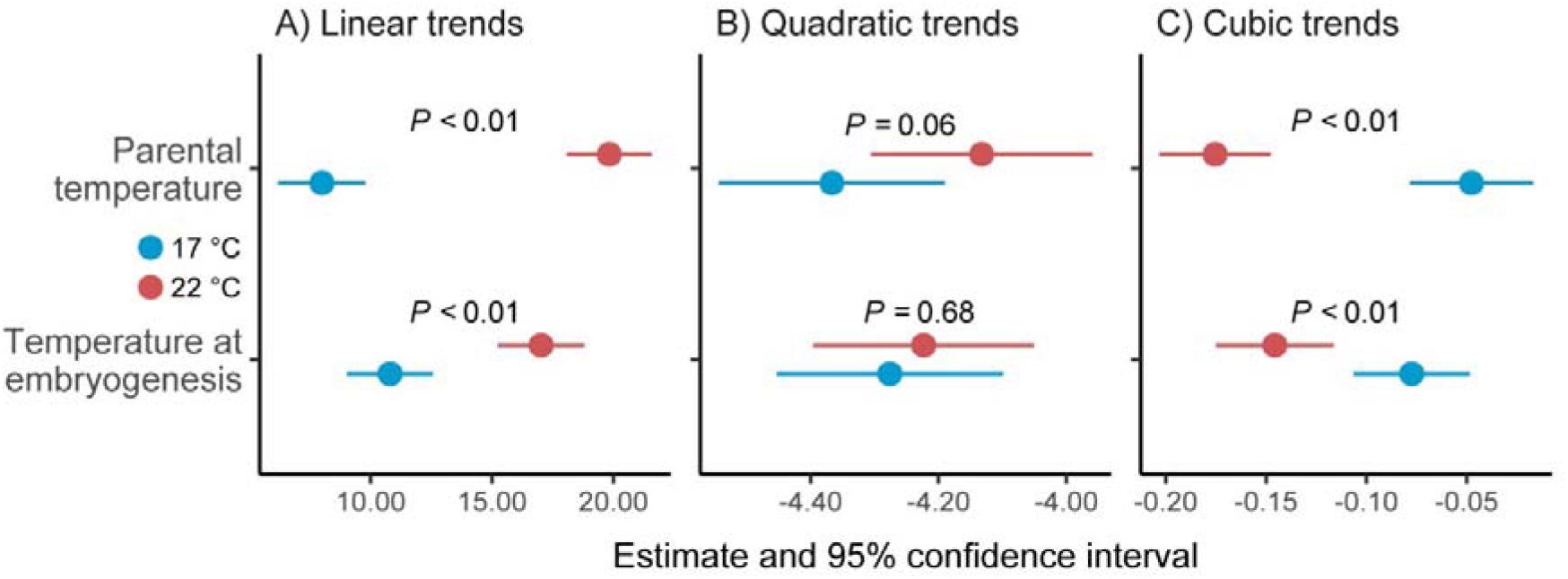
Independent effects of parental temperature and temperature at embryogenesis (17 °C in blue *versus* 22 °C in red) on A) linear, B) quadratic, and C) cubic trends estimated for survival curves in Figure 4. Trends for parental temperatures are averaged across temperatures at embryogenesis, and *vice versa.* All estimates (points) and 95% confidence intervals exclude zero and are multiplied by 100 to aid visualization of different scales. Survival was unaffected by temperature at fertilization.

Temperature at fertilization did not affect mean survival, alone or in combination with any other acclimation temperature (Table 1).

### Estimates and confidence intervals of curve descriptors

As suggested by survival curves above, estimates and confidence intervals for curve descriptors revealed stronger responses to parental temperature than to temperature at embryogenesis.

Independent of temperature at embryogenesis, maximal survival of larval development was significantly higher when parents acclimated at 22 °C than when they acclimated at 17 °C (Figure 6A). Consistent with thermodynamic effects on survival, moreover (Figure 1C), the significant increase in maximal survival was accompanied by a significant increase in the thermal optimum for survival, which was 1.2 °C higher (and confidence intervals did not overlap) when parents acclimated at 22 °C than when they acclimated at 17 °C (Figure 6B).

**Figure 6.**
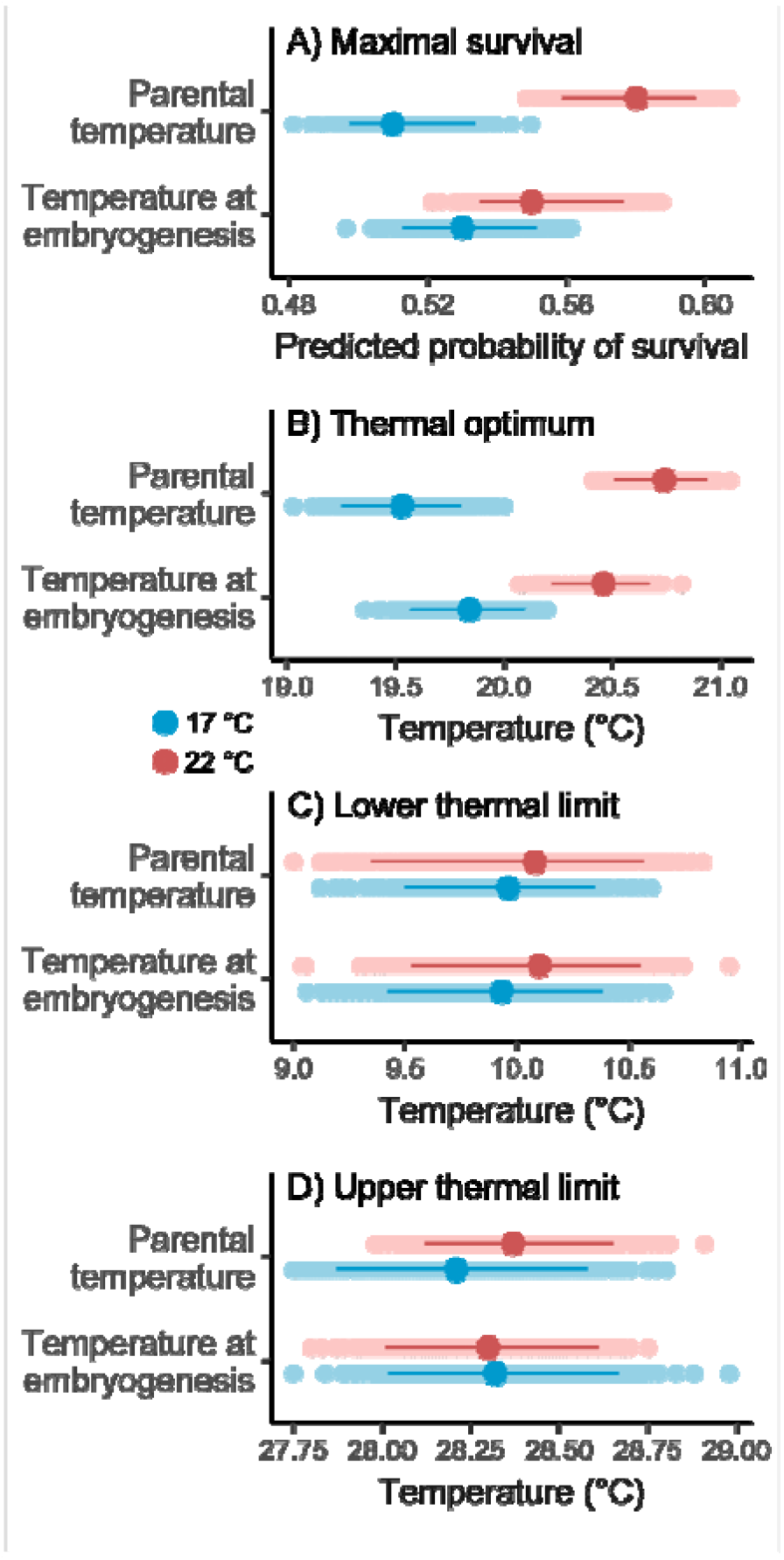
Effects of parental temperature and temperature at embryogenesis (17 °C in blue *versus* 22 °C in red) on descriptors of survival curves in Figure 4: A) maximal survival, B) thermal optimum, and C-D) thermal limits. Descriptors for parental temperatures are averaged across temperatures at embryogenesis, and *vice versa.* Estimates (solid points) and 95% confidence intervals are based on 1,000 bootstrap replicates (transparent points) of the binomial mixed-effects regression used to model curves (see bootstrapping details in text). Survival was unaffected by temperature at fertilization.

Independent of parental temperature, higher temperature at embryogenesis also increased the thermal optimum for survival, which was 0.6 °C higher when embryos developed at 22 °C than when they developed at 17 °C (Figure 6B). Although the increase in optimum again coincided with an increase in maximal survival, this increase was not significant (Figure 6A). Hence, the effects of temperature at embryogenesis on curve descriptors were in the directions predicted for thermodynamic effects (Figure 1C), but inseparable statistically from the predictions for beneficial acclimation (Figure 1A).

Thermal limits for survival were unresponsive to prior acclimation temperatures (Figure 6C-D), as were thermal breadth and thermal tolerance (not presented).

## Discussion

Understanding the fitness consequences of thermal history is necessary to predict organismal responses to global warming, especially for ectotherms with distinct life stages that can differ in thermal sensitivity, acclimate to different thermal environments, and accrue responses within and between generations (Williams *et al.* 2016; Kellermann, van Heerwaarden & Sgrò 2017; Donelson *et al.* 2018). While acclimation is often hypothesized to benefit organisms by helping them (or their offspring) to compensate for negative impacts of environmental change (Leroi, Bennett & Lenski 1994; Uller, Nakagawa & English 2013), equivocal support for this hypothesis (Huey *et al.* 1999; Sgrò, Terblanche & Hoffmann 2016; Sánchez□Tójar *et al.* 2020) highlights the need to assess alternatives. Here in *Galeolaria,* an aquatic ectotherm whose early planktonic stages (gametes, embryos, and larvae) are especially vulnerable to warming (Walsh *et al.* 2019; Byrne *et al.* 2020; Dahlke *et al.* 2020), we show that thermal survival curves are most responsive to parental acclimation followed by acclimation at embryogenesis, but are buffered against acclimation at fertilization. Moreover, curves respond independently to acclimation within and between generations, and respond largely as predicted by the warmer-is-better hypothesis, despite the semblance of beneficial acclimation after successive acclimations to warmer temperature. Our results demonstrate the varied nature of thermal acclimation, and the importance of considering how acclimation responses accumulate across complex life cycles when predicting the impacts of warming.

Drawing on thermodynamic principles, the warmer-is-better hypothesis argues that adapting to warmer temperatures confers greater bioenergetic capacity than adapting to cooler temperatures (Figure 1C; Angilletta, Huey & Frazier 2010), and is invoked to explain why maximal performance is often higher in organisms with high thermal optima than those with lower optima (Sørensen *et al.* 2018; Treasure & Chown 2019; Buckley & Kingsolver 2021). To our knowledge, such a response to parental acclimation has not yet been documented, despite ongoing scrutiny of parental effects and debate over their adaptive value (Uller, Nakagawa & English 2013; Donelson *et al.* 2018; Sánchez□Tójar *et al.* 2020). This could reflect the long tradition of designing experiments to test beneficial acclimation by assessing offspring fitness in environments either matched or mismatched to that of parents (as synthesised in Uller, Nakagawa & English 2013), without testing other hypotheses by assessing joint shifts in features of thermal performance curves (Huey *et al.* 1999; Sørensen *et al.* 2018). Mechanisms underlying the transfer of acclimation effects from parents to offspring remain contentious (McGuigan, Hoffmann & Sgrò 2021), but may involve packaging cellular defenses such as stress-response proteins or hormones into developing gametes before fertilization (Burton & Metcalfe 2014; Lockwood, Julick & Montooth 2017; Gulyas & Powell 2019). Such defenses are upregulated at warmer temperatures, give frontline protection against thermal stress, and are not synthesized by mature gametes or early embryos (Feder & Hofmann 1999; Sørensen, Kristensen & Loeschcke 2003; Hamdoun & Epel 2007), suggesting a plausible pathway for parental acclimation to produce warmer-is-better effects in offspring. This pathway may also explain the buffering of survival in *Galeolaria* against acclimation at fertilization, which (consistent with Figure 1D) our analyses could ignore without loss of fit, By contrast, acclimation of embryos had more ambiguous effects on thermal survival curves. Statistically, at least, larvae compensated for warmer temperature at embryogenesis by increasing their thermal optimum at no cost maximal survival, as predicted for the beneficial acclimation hypothesis (Figure 1A; Leroi, Bennett & Lenski 1994; Sørensen *et al.* 2018). However, the increase was weaker than that induced by parental acclimation to the same temperature, and accompanied by a correspondingly weak (non-significant) increase in maximal survival, suggesting that joint responses to acclimation at this life stage are still in the general direction of the warmer-is-better hypothesis. Such a result could again point to the involvement of stress-response proteins, which often go unexpressed until later in embryogenesis because their overexpression during early cleavages can inhibit cell division and signaling (Feder & Hofmann 1999; Hamdoun & Epel 2007). Planktonic embryos are therefore thought to evolve rapid development to speed progression through risky stages with less protection against stress (Strathmann, Staver & Hoffman 2002). The shortened window of exposure, and non-expression of cellular defenses for part of it, might then give embryos less scope than parents to respond to acclimation and transfer responses to future life stages. Alternatively, cumulative damage from successive bouts of acclimation might counter any fitness gains from thermodynamic effects (Williams *et al.* 2016; Buckley & Kingsolver 2021), as inferred by enhanced survival of *Galeolaria* larvae (independent of test temperature) when parental acclimation at 22 °C was followed by embryogenesis at 17 °C, but not embryogenesis at 22 °C (Figure 3). If so, the semblance of beneficial acclimation could emerge as a by-product of thermodynamics, thereby giving such effects a potential role in adaptive responses to climate change (Williams *et al.* 2016; Sørensen *et al.* 2018).

The relative impacts of acclimation within generations *versus* between generations here go against meta-analytic findings that parental environmental effects do not significantly buffer offspring survival against stress (Uller, Nakagawa & English 2013; Sánchez□Tójar *et al.* 2020), and are weaker than offspring environmental effects (though these may relate to test environments only; Uller, Nakagawa & English 2013). One reason may be that such analyses synthesize a broad range of environmental factors with differing effect sizes. However, targeted reviews of offspring responses to parental temperature report similarly equivocal results (Sgrò, Terblanche & Hoffmann 2016; Donelson *et al.* 2018). Another reason may be that the life history characteristics of aquatic organisms, combined with higher thermal inertia (and hence higher predictability of temperature cues) in water than on land, offer more scope for mechanisms of temperature compensation to evolve (Sørensen *et al.* 2018). In external fertilizers like *Galeolaria,* for example, high mortality at early life stages that disperse passively in currents can lead to strong coupling of physical and evolutionary processes, while decoupling the environments of early stages and adults (Lotterhos, Albecker & Trussell 2021). As such, temperature may vary more between generations than within them (at least for the stages considered here), favoring stronger responses to parental acclimation than to acclimation in early development (Angilletta 2009; Le Roy & Seebacher 2018). While further tests are needed, a recent review citing beneficial responses to transgenerational acclimation in 47% of studies on aquatic invertebrates, compared to 26% of studies on terrestrial ones, offers some qualified support for this idea (Donelson *et al.* 2018; see also Byrne *et al.* 2020).

To our knowledge, our study remains one of only a few to explicitly dissect the effects of parental and within-generation acclimation on thermal performance (Huey *et al.* 1995; Zamudio, Huey & Crill 1995; see also Le Roy & Seebacher 2018), though others have done so for within-generation acclimation alone (Deere & Chown 2006; Treasure & Chown 2019). More often, as noted above, parental acclimation temperatures are crossed factorially with offspring test temperatures in ways that can neither distinguish among competing hypotheses of acclimation, nor assess cumulative responses to it (but see Steigenga & Fischer 2007). Consequently, we lack a clear picture of how such responses accrue within generations and from one generation to the next (Williams *et al.* 2016; Buckley & Kingsolver 2021). In other work on *Galeolaria,* for instance, larval survival at warmer temperature was enhanced by warm acclimation of parents (Chirgwin *et al.* 2018), but reduced by warm acclimation of gametes (Chirgwin, Connallon & Monro 2021), while warm acclimation of embryos increased the optimum temperature for survival by ~2 °C (Rebolledo, Sgrò & Monro 2021). However, temperatures at postzygotic stages were conflated in the first two studies, and parents were unacclimated in the latter two. Results here now suggest that our interpretation of acclimation responses, and ensuing predictions of climate vulnerability, may change if such responses are considered in aggregate across complex life cycles.

Overall, thermal acclimation in *Galeolaria* differs within and between generations in ways that support differing hypotheses about its impacts on thermal sensitivity, and suggest the transfer of thermodynamic (warmer-is-better) effects from parents to offspring. Warm acclimation of parents, and to less extent embryos, enhanced larval survival at warmer temperatures, without affecting thermal limits or breadth, to thereby expand the area under the curve relating survival to temperature. The extent to which this may buffer vulnerable planktonic stages against warming, however, is unclear. Our simulated 5 °C of warming exceeds projected regional levels of ~2-4 °C by the end of the century under a high emissions pathway (RCP8.5 relative to 1980-1999; Lough, Sen Gupta & Hobday 2012), but increased the thermal optimum for survival by only ~2 °C on aggregate. Acclimation might thus compensate for warming below worst-case scenarios, but not extreme climate events such as heatwaves, which are already warming regional waters by up to 5 °C and for periods longer than *Galeolaria*’s life cycle (Oliver *et al.* 2017; Babcock *et al.* 2019). Future work should therefore assess acclimation to more extreme temperatures than studied here, to find the range at which it can no longer prevent the accumulation of damage and homeostasis is disrupted to the point of mortality (Ørsted, Jørgensen & Overgaard 2022). Another key step will be to assess potential trade-offs, for example, between parents’ investment in their own defenses against warming *versus* those of offspring (Waite & Sorte 2022). Nevertheless, our results imply that gradual warming may benefit ectotherms not yet living at their thermal limits if acclimation induces thermodynamic effects that enhance survival, or prove detrimental if survival declines because mechanisms of temperature compensation are disrupted. Our study therefore demonstrates the importance of understanding how acclimation responses aggregate across complex life cycles when predicting the impacts of warming.

## Acknowledgements

We thank Cristóbal Gallegos Sánchez and Emily Belcher for help in collecting specimens, Craig White for contributing equipment, Vanessa Kellermann and Mads Schou for contributing R code, and Fisheries Victoria for collection permits.

## Author contributions

A.P.R., K.M. and C.M.S. conceived the ideas and designed methodology; A.P.R. collected the data; A.P.R. and K.M. analyzed the data, created the graphics and drafted the manuscript; all authors contributed to revisions and gave final approval for publication.

## Data availability statement

Data will be available upon acceptance of the manuscript.

